# Statistically robust methylation calling for whole-transcriptome bisulfite sequencing reveals distinct methylation patterns for mouse RNAs

**DOI:** 10.1101/130419

**Authors:** Carine Legrand, Francesca Tuorto, Mark Hartmann, Reinhard Liebers, Dominik Jacob, Mark Helm, Frank Lyko

## Abstract

Cytosine-5 RNA methylation plays an important role in several biologically and pathologically relevant processes. However, owing to methodological limitations, the transcriptome-wide distribution of this mark has remained largely unknown. We previously established RNA bisulfite sequencing as a method for the analysis of RNA cytosine-5 methylation patterns at single-base resolution. More recently, next-generation sequencing has provided opportunities to establish transcriptome-wide maps of this modification. Here we present a computational approach that integrates tailored filtering and data-driven statistical modeling to eliminate many of the artifacts that are known to be associated with bisulfite sequencing. Using RNAs from mouse embryonic stem cells we performed a comprehensive methylation analysis of mouse tRNAs, rRNAs and mRNAs. Our approach identified all known methylation marks in tRNA and two previously unknown but evolutionary conserved marks in 28S rRNA. In addition, mRNAs were found to be very sparsely methylated or not methylated at all. Finally, the tRNA-specific activity of the DNMT2 methyltransferase could be resolved at single-base resolution, which provided important further validation. Our approach can be used to profile cytosine-5 RNA methylation patterns in many experimental contexts and will be important for understanding the function of cytosine-5 RNA methylation in RNA biology and in human disease.

## Introduction

5-methylcytosine (m5C) is the longest-known and best-understood epigenetic modification of DNA (Jones 2012). The genome-wide analysis of m5C patterns has greatly aided to our understanding of epigenetic gene regulation (Bock et al. 2010). Changes in DNA methylation patterns have been found to underpin organismal development and cellular differentiation (Smith and Meissner 2013) and also provide valuable biomarkers for the detection of human diseases, including cancer (Heyn and Esteller 2012).

m5C also represents a well-known modification of RNA (Motorin et al. 2010). In comparison to DNA modifications, RNA modifications are substantially more diverse and complex, but their functional significance is only beginning to be elucidated (Gilbert et al. 2016; Tuorto and Lyko 2016). RNA modifications are particularly enriched in tRNAs, where they are often linked to translational regulation (Agris 2008). In this context, it has been shown that m5C modification of tRNA plays an important role in tRNA stability and in the regulation of translational fidelity (Schaefer et al. 2010; Tuorto et al. 2012; Blanco et al. 2014; Tuorto et al. 2015). Furthermore, m5C is also a widely conserved modification of rRNA, where it is implied in the quality control of ribosome biogenesis (Sharma et al. 2013; Bourgeois et al. 2015; Schosserer et al. 2015). These processes have been associated with a variety of human diseases (Blanco and Frye 2014).

m5C in RNA can be reliably detected using radioactive labeling and thin-layer chromatography (Hengesbach et al. 2008). However, this method only allows for an indirect quantification of global methylation levels. In comparison, high performance liquid chromatography coupled to mass spectrometry (LC-MS) is more accurate and also allows the analysis of larger sample numbers (Thuring et al. 2016), but currently does not provide any information about the sequence context of the methylation marks. Several methods for the mapping of RNA cytosine methylation marks have been proposed (Hussain et al. 2013a; Li et al. 2016), however, they are usually based on indirect detection and thus represent approximations of the actual distribution. Direct mapping of RNA m5C marks in their native sequence context is currently only provided by RNA bisulfite sequencing (Schaefer et al. 2009). Bisulfite sequencing is based on the selective deamination of unmethylated cytosines, thus converting unprotected cytosines to uracils, followed by sequencing-based detection of methylation-related sequence polymorphisms (Clark et al. 1994). RNA bisulfite sequencing can accurately identify the presence of selected known methylation marks in tRNA and rRNA and has proven to be very useful for the molecular characterization of RNA cytosine-5 methyltransferases (Schaefer et al. 2009).

A few studies have also utilized RNA bisulfite sequencing to map RNA m5C marks at the transcriptome level and found evidence for the presence of m5C in mammalian mRNAs and non-coding RNAs (Squires et al. 2012; Hussain et al. 2013b; Khoddami and Cairns 2013; Amort et al. 2017). While it has been suggested that the human coding and non-coding transcriptome contains up to 10,000 methylation sites (Squires et al. 2012; Amort et al. 2017), the function of these marks has remained elusive. Furthermore, the available studies could not define a common set of substrate mRNAs or consensus methylation target sequences, which raised the possibility that some of the results were influenced by incomplete deamination, secondary structures or other confounding factors that are known to affect bisulfite sequencing (Supplemental Table S1). It is also possible that methylation calling was influenced by insufficient statistical stringency.

Standard analytical pipelines for whole-transcriptome bisulfite sequencing datasets are currently not available. Furthermore, available tools do not sufficiently address the need for statistical approaches in the elimination of stochastic artifacts. Our approach addresses these issues and identifies profoundly different methylation patterns for mRNA, rRNA and tRNA.

## Results

Whole-transcriptome bisulfite sequencing libraries were prepared by separating total RNA samples into small (<200 nt) and long (>200 nt) RNA fractions. Depending on whether ribosomal RNA was also examined, an rRNA depletion step was included or not. The long fraction was fragmented to a size distribution appropriate for Illumina sequencing. Both fractions were DNase digested, bisulfite converted and end-repaired prior to cDNA library preparation for deep sequencing (Fig. 1A; Supplemental Methods). Sequenced reads were aligned with BSMAP and subjected to initial quality control (read length ≥25 nt, aligned uniquely, forward and reverse reads at the same location). A closer examination of candidate methylation sites revealed many obvious false positive sites that were related to inefficient conversion (i.e. tracts of ≥3 consecutive unconverted Cs) or misalignment (Supplemental Fig. S1). We therefore implemented filters for the removal of these artifacts (see Supplemental Methods for details). For methylation calling, a Poisson distribution was fit to each sample, and non-conversion p-values were calculated. Finally, available replicates were joined and a combined non-conversion p-value was calculated (Fig. 1B; Methods).

**Figure 1.**
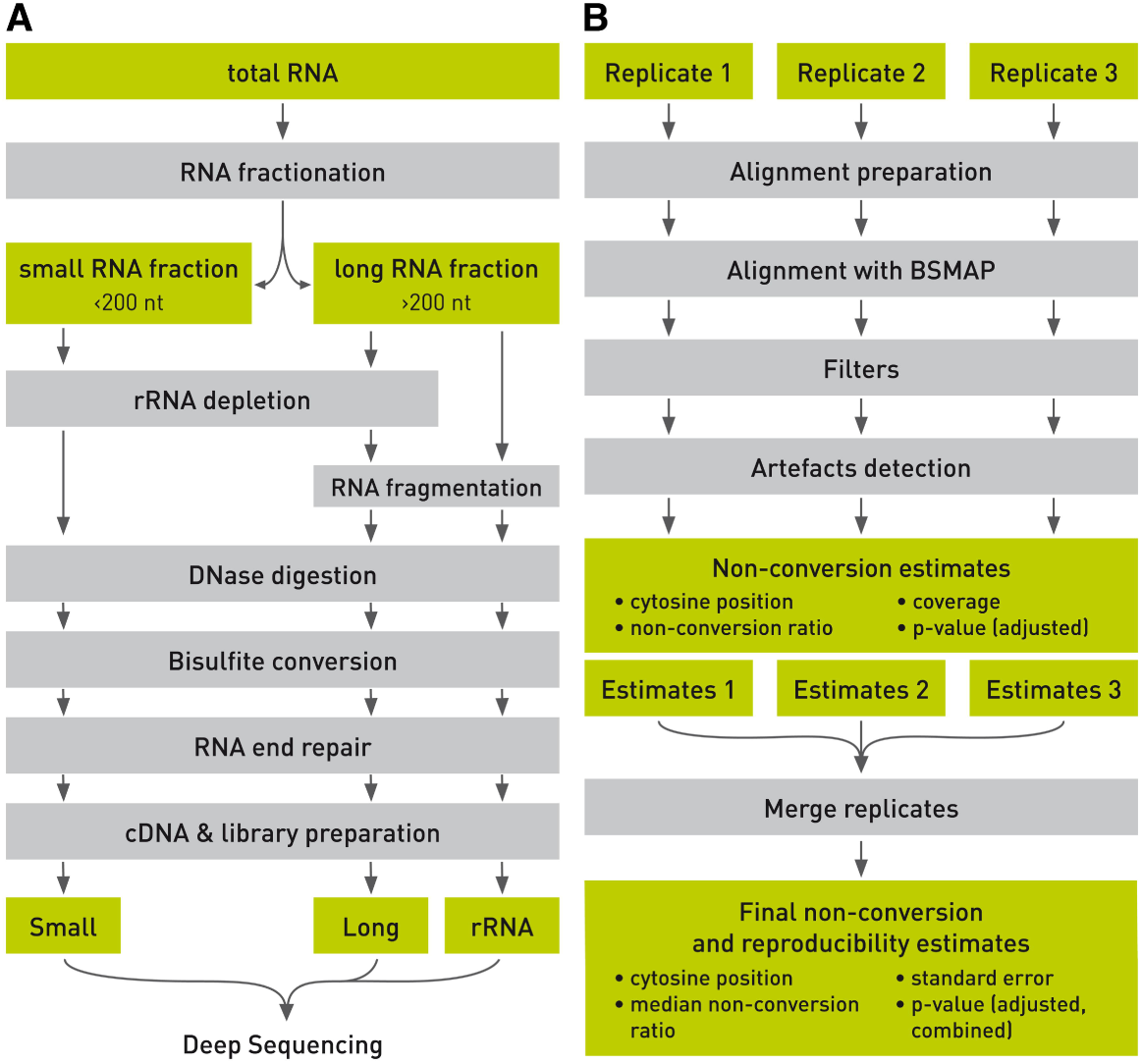
Schematic outline of whole-transcriptome bisulfite sequencing. The illustration shows key steps of library preparation (*A*) and data analysis (*B*).

For the present study, we sequenced and analyzed 3 replicates of bisulfite-converted libraries from mouse embryonic stem (ES) cells and various genotypes. 19,198,431 to 84,435,380 read pairs were available after sequencing of each sample (Supplemental Table S2). Bisulfite conversion rates were determined from unmethylated regions of rRNA and ranged from 98.4% to 98.6% (Supplemental Table S2).

Initial data analysis using the mRNA dataset from a single wild type replicate showed that the vast majority of cytosines had conversion ratios larger than 90% (Fig. 2A). To further analyze the non-converted cytosine reads, we obtained the Poisson parameter λ; from non-converted cytosine counts for each coverage bin between 10x and 1,200x (with average and median coverages of this subset being 44x and 20x, respectively) and calculated the median λ_*p*_/*coverage* as 0.0164 with 95% confidence interval [0.0164; 0.0166] (Fig. 2B, Supplemental Table S3). Not surprisingly, (1 - λ_*p*_/*coverage*) is close to the deamination rate for this sample, which we calculated at 98.3% (Supplemental Table S2). Similarly, the ratios λ_*p*_/*coverage* were obtained for all sequencing libraries (Supplemental Table S3) and showed only minor differences, in agreement with minor variations in bisulfite deamination efficiencies (Supplemental Table S2).

**Figure 2.**
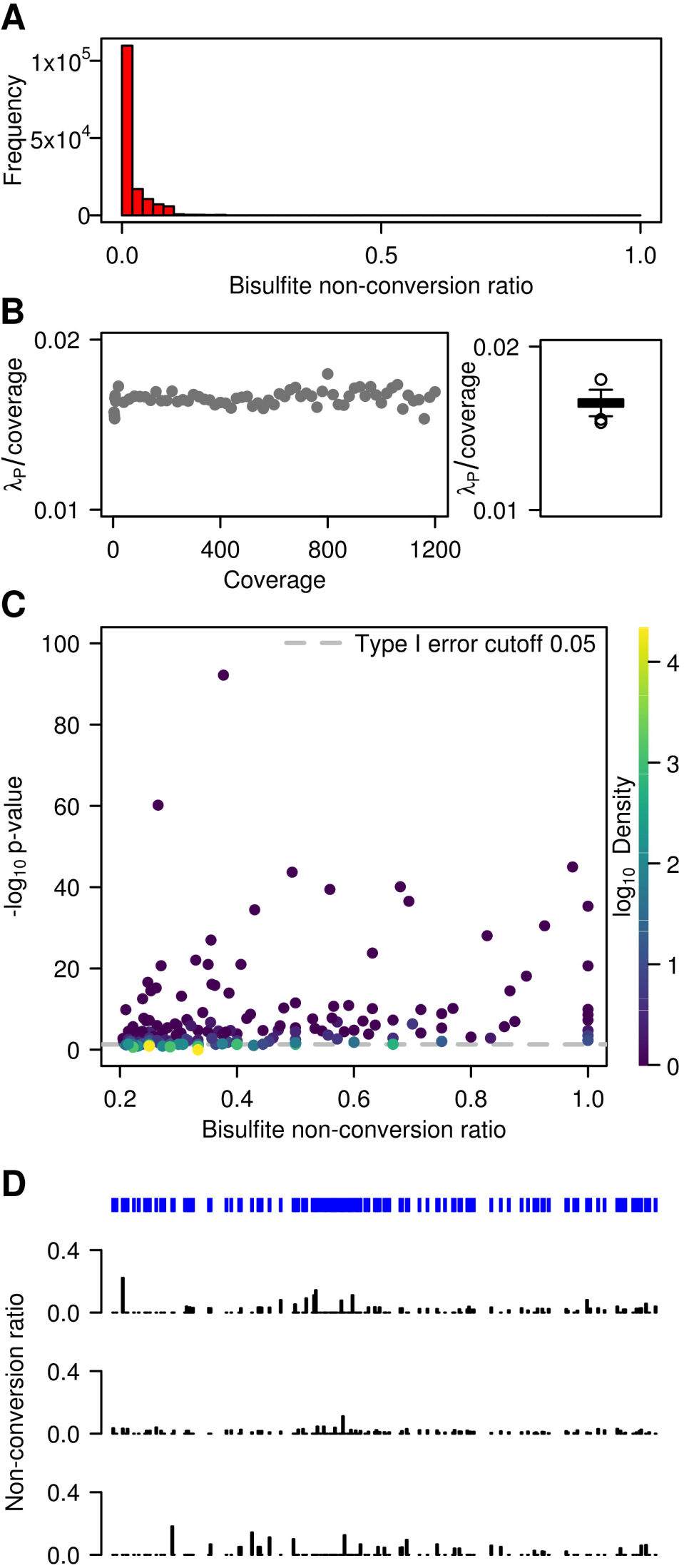
WTBS of mouse ES cell mRNA. (*A*) Frequency of bisulfite non-conversion (resp. methylation) ratios in mRNA. (*B*) Estimate Poisson rate λ_*p*_/*coverage* as a function of coverage, and corresponding boxplot. (*C*) Bisulfite non-conversion ratio (x-axis) and - log of methylation p-values (y-axis) for 9,959 cytosines passing the 0.2 ratio threshold. Color indicates density (count of points per symbol area). (*D*) Cytosine position and methylation tracks for a representative mRNA transcript (NM_007984) in three replicates.

In subsequent steps, we compared non-conversion rates with the underlying distribution for each cytosine site where the non-conversion ratio is higher than λ_*p*_/*coverage*. Out of 3,338,384 cytosines, 53,510 had a non-conversion ratio higher or equal to 0.2 (Fig. 2C), consistent with earlier findings in human cell lines (Squires et al. 2012). However, only 266 out of 53,510 cytosines achieved statistical significance (p<0.05) after Benjamini-Hochberg correction for multiple testing (Fig. 2C). Many significant sites exhibited a low non-conversion ratio of 0.2. Conversely, some cytosines with higher non-conversion ratios did not pass a 0.05 significance threshold (Fig. 2C). The large amount of non-significant p-values demonstrates the requirement for statistical approaches in the analysis of whole-transcriptome bisulfite sequencing datasets. This notion was confirmed when we analyzed non-conversion tracks for all 3 independent replicates (see representative example in Fig. 2D). In spite of the considerable overall deamination efficiency, all samples showed a persistent background of residual non-conversion as well as individual sites with particularly reduced conversion rates (Fig. 2D). However, these patterns were usually not conserved between the 3 replicates, indicating that they are the result of non-reproducible methylation or random incomplete bisulfite deamination.

In subsequent analyses, we therefore examined the intersection of the 3 replicates (i.e., sites with a non-conversion rate consistently above the library non-conversion rate shown in Supplemental Table S2) to reliably identify methylated cytosines in mRNAs. Out of the 2,105,654 cytosines that had sufficient coverage in all 3 replicates, 56,940 had a bisulfite non-conversion ratio larger than λ_*p*_/*coverage*. Remarkably, only 745 sites were significant (adjusted p-value <0.05, Fig. 3A). Out of these candidates, only a small fraction combined high statistical significance with high methylation ratios (Fig. 3A; Supplemental Table S3). In order to assess the type I error of the method, we also evaluated the proportion of false positives resulting from the Poisson test by a simulation of several instances of 1,000,000 stochastically non-converted cytosines. This allowed us to reconstitute non-conversion ratios (Fig. 3B) and to calculate type I error estimates (Supplemental Fig. S2). Simulated non-conversion ratios were remarkably close to the experimental data (Fig. 3B), thus confirming that our simulations are realistic, and type I error was identical or lower than the significance level of 0.05 (Supplemental Fig. S2). Simulations of 10,000 cytosines methylated at various levels among 3,000,000 cytosines provided statistical power estimates, using either Benjamini-Hochberg adjustment (Fig. 3C) or Independent Hypothesis Weighting (Supplemental Methods, Supplemental Fig. S2) for multiple testing. The results showed that our analysis had the ability to reliably (statistical power = 99.9±0.04%) detect sites with as low as 20% non-conversion at a coverage of >20. A coverage of >20 was obtained for >50% of the cytosine residues in our sequencing datasets (Fig. 3D), thus demonstrating sufficient statistical power for a meaningful data analysis.

**Figure 3.**
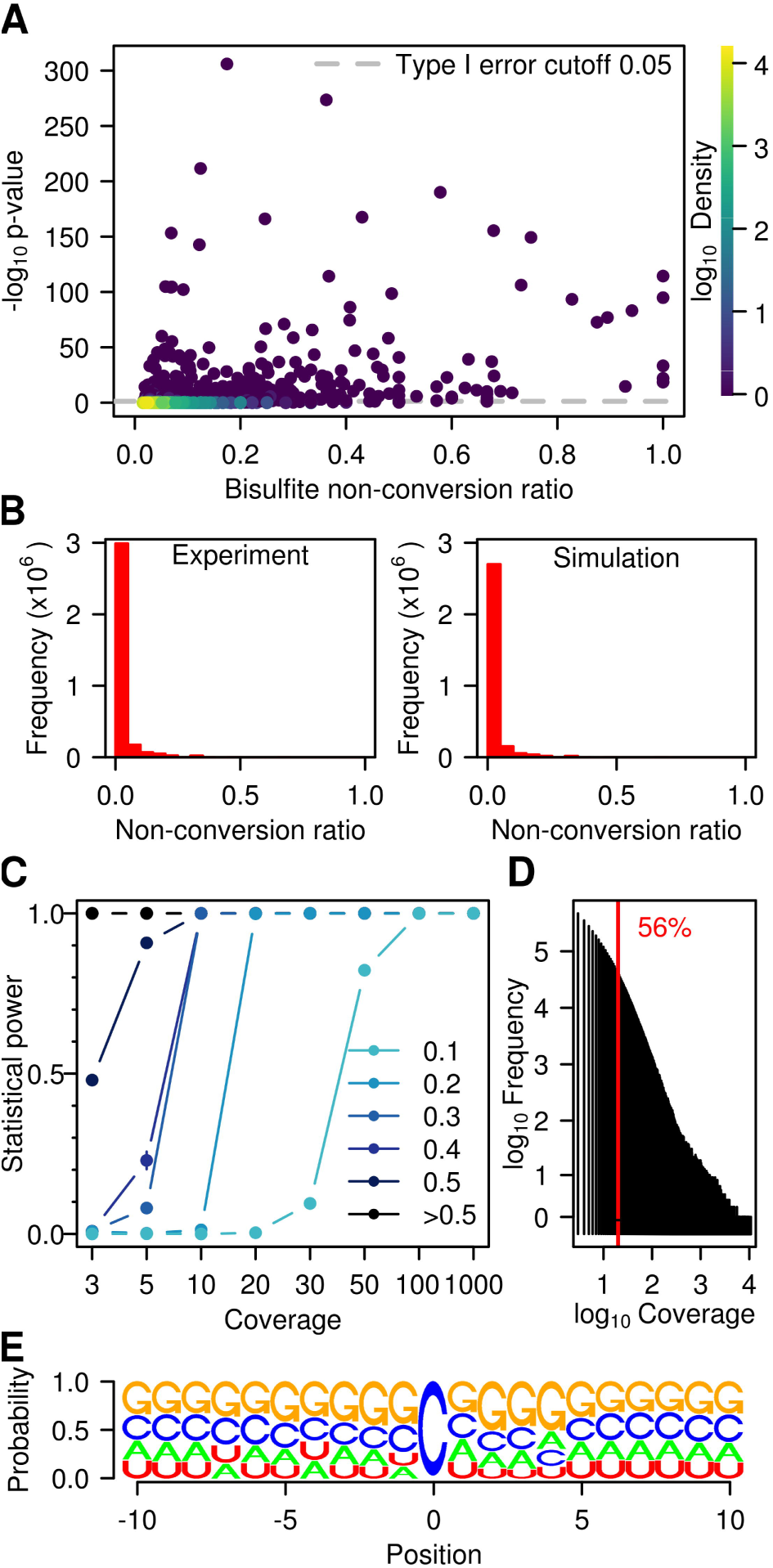
Statistical analysis of WTBS datasets. (*A*) Bisulfite non-conversion ratio (x-axis) and - log of methylation p-values (y-axis) for 7,445 cytosines with ratio > λ_*p*_/*coverage* that are common to the three wild type replicates. Color indicates density (count of points per symbol area). (B) Number of cytosines and non-conversion ratios, as determined experimentally (left panel) or by simulation (right panel). (C) Statistical power stratified by coverage and by non-conversion ratio. (D) Histogram of coverage in a representative (Wt1L) sequencing data set. The red line indicates a coverage of 20. (*E*) Logo plot for all cytosines with ratio >0.1 and significant p-value (<0.05) in at least one sample.

However, even after statistical analysis, several candidate sites remained that could not be considered bona fide methylation sites. For example, site *C153* within mRNA NM_001199350 had a median non-conversion ratio of 1.0 and an adjusted p-value of 6.6 x 10^-84^, but actually presented a C>T sequence polymorphism at this specific position. As a consequence, a read containing the T variant multiply aligned to both the NM_001199350 sequence and to its T-containing variant, and was therefore discarded. However, C-containing reads would align uniquely and were therefore maintained in the analysis, resulting in an overestimation of the methylation ratio. To further address the reproducibilty of the remaining candidate methylation sites, we selected 10 candidates with the lowest adjusted p-values and 4 additional candidates with methylation level close to 1.0 and significant adjusted p-values in a subset of samples (Supplemental Table S4) for an amplicon-based re-sequencing approach. The results showed that 4 out of 14 analyzed cytosines were unmethylated (Supplemental Fig. S3; Supplemental Table S4). This again suggests that the very low number of candidate methylation sites identified in our analysis contains a certain amount of false positives. In agreement with this notion, candidate methylation sites also failed to reveal any pattern specificity, as we could not identify any clear enrichment for specific sequence contexts (Fig. 3E).

The prevalence of m5C in mRNA was further analyzed by LC-MS/MS. mRNA was enriched from total RNA by two consecutive rounds of polyA selection, followed by smallRNA depletion and two consecutive rounds of rRNA depletion (Fig. 4A). Samples were taken at each step and analyzed by RNA-seq for sample composition and by LC-MS/MS for base modifications. The sequencing results showed that the enrichment protocol resulted in a strong increase of mRNA reads (Fig. 4B). However, a significant (7.1 %) fraction of rRNA reads remained after the final step (Fig. 4B), probably resulting from ineffective rRNA fragment depletion. LC-MS/MS analysis of all samples demonstrated a strong reduction of m5C (Fig. 4C) that closely corresponds to the rRNA depletion observed by sequencing. Of note, we also failed to detect any evidence for the presence of the oxidated derivative of m5C, 5-hydroxymethylcytosine in any of our samples (Fig. 4C). Very similar results were obtained in parallel analyses of *Drosophila* S2 cell RNA samples (Supplemental Fig. S4). Based on a detection limit of 1 fmol (Supplemental Fig. S5), this finding corresponds to a maximum hm5C content of 50 ppm (0.005%) per C residue. Our results thus contrast the antibody-based detection of 5-hydroxymethylcytosine as a prevalent mRNA modification in *Drosophila* (Delatte et al. 2016). Furthermore, our results also suggest that mRNAs are very sparsely methylated or not methylated at all.

**Figure 4.**
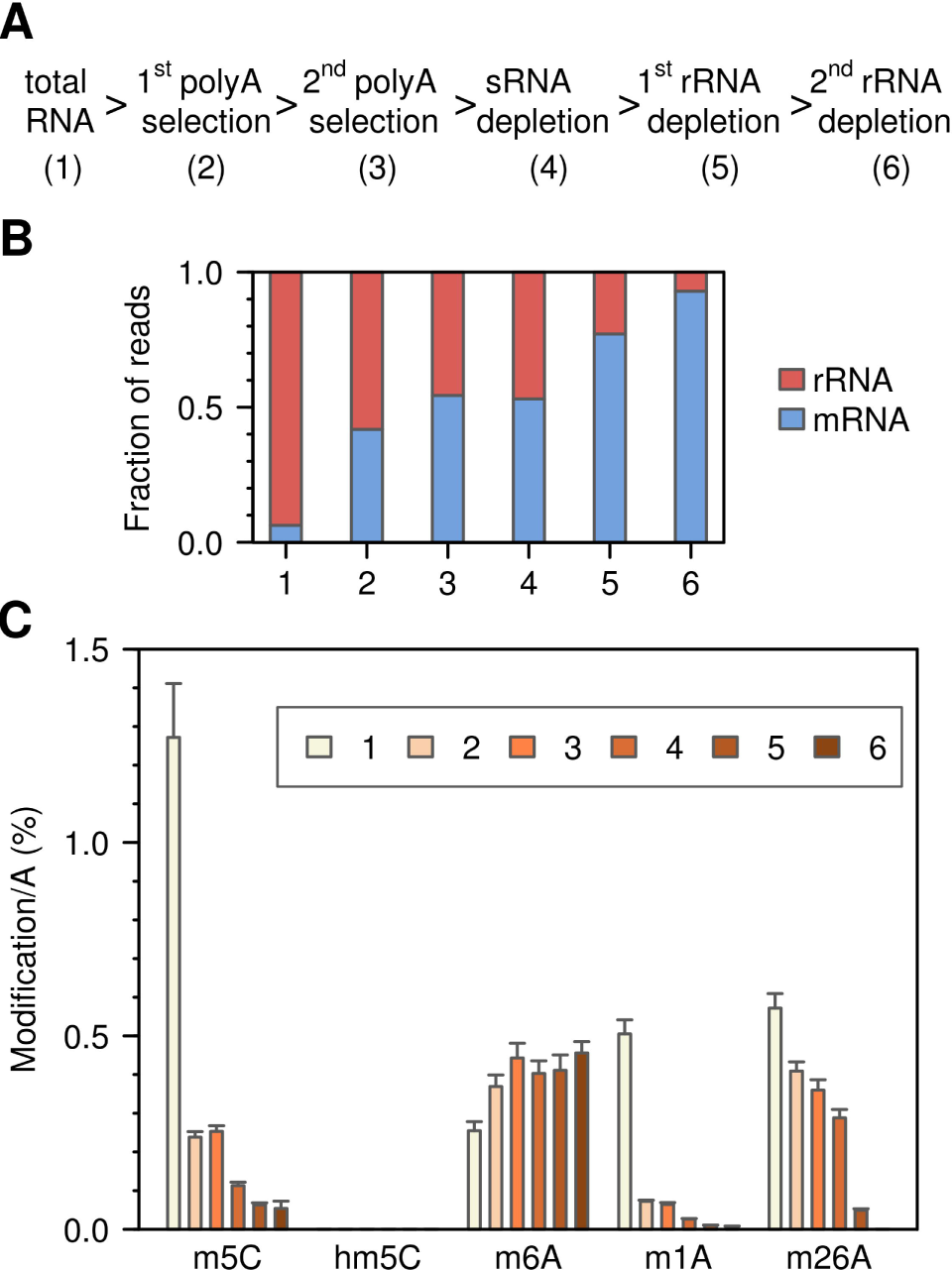
LC-MS/MS analysis of RNA samples from mouse ES cells that were subjected to multi-step mRNA enrichment. (A) Basic outline of the mRNA enrichment protocol. (B) Relative amounts of mRNA and rRNA, as determined by RNA-seq. The proportion of tRNA reads was <1 % for all samples. (C) Modification analysis of m5C, hm5C, m26A, m6A and m1A content, relative to A content.

While the cytosine-5 methylation status of mRNAs is discussed controversially, rRNAs and tRNAs have long been known to carry defined methylation marks (Machnicka et al. 2013). We therefore compared the methylation frequencies between these three types of RNA. In mRNA, only 20 cytosines out of 43,410 had a significant bisulfite non-conversion ratio larger than 0.1, which corresponds to a hypothetical mRNA methylation level of 0.05% (Fig. 5A). In contrast, methylation levels appeared to be substantially higher in rRNA (1.3%) and in tRNA (8.6%), which is consistent with the known prevalence of m5C in these RNAs (Fig. 5A). Similarly, when we systematically evaluated the reproducibility of the non-conversion ratio for all candidate methylation sites, standard errors appeared substantially higher for mRNA than for rRNA and tRNA (Fig. 5B). This finding again suggests that tRNAs and rRNAs are methylated differently from mRNAs.

**Figure 5.**
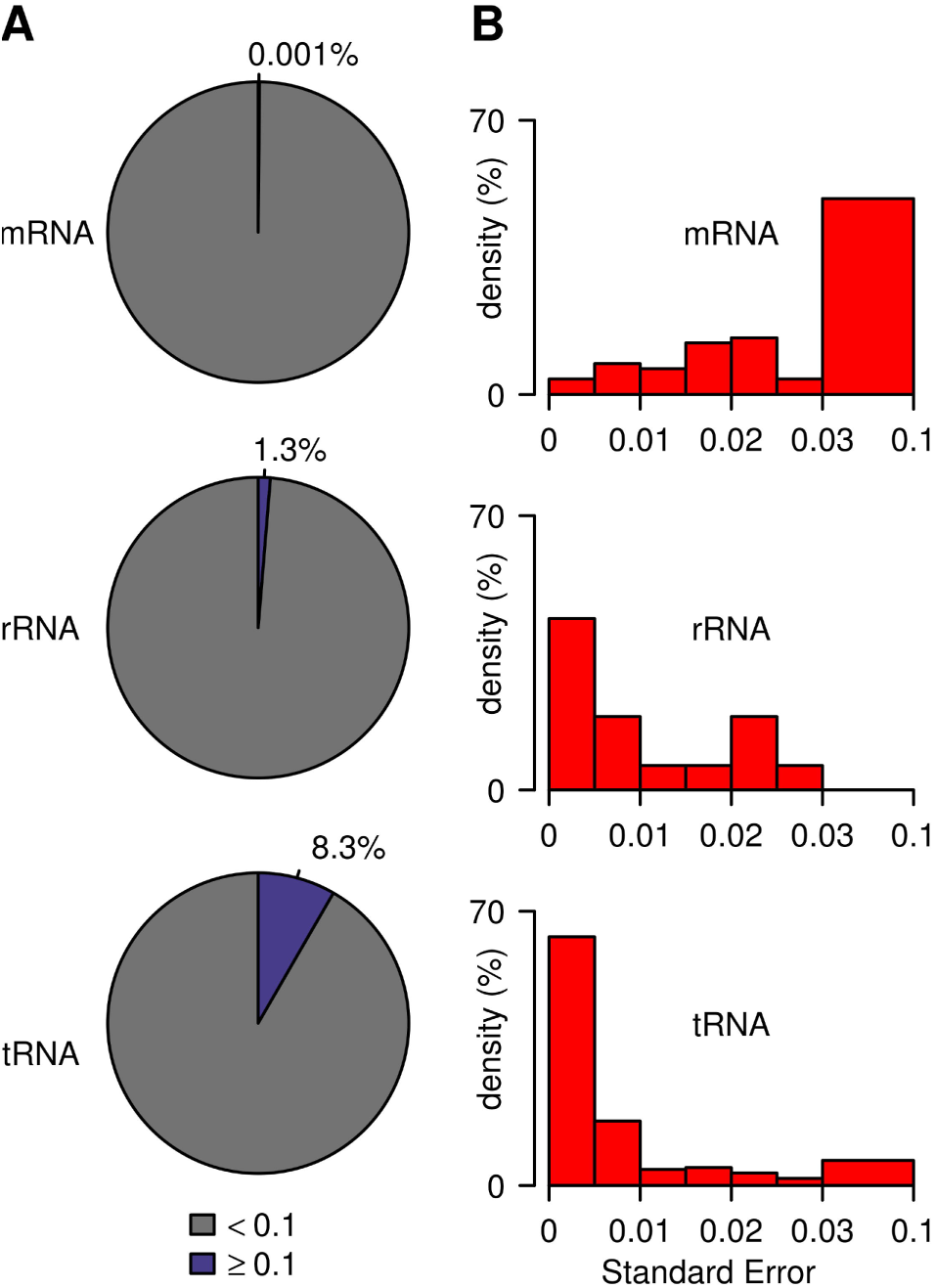
Methylation frequencies and reproducibility in mRNA, rRNA and tRNA. (*A*) Proportion of non-converted cytosines (coverage ≥ 20). (*B*) Standard error (coverage ≥ 20 and non-conversion ratio ≥ 0.1). Values are based on triplicate wild type datasets for mRNA and tRNA, and duplicate datasets for rRNA.

Finally, we also investigated the effect of specific enzymes on RNA m5C patterns. To this end, we further developed our computational pipeline to provide a standard analysis of specific types of RNA and to compare methylation patterns between genotypes. As a proof of principle, we determined the effects of the presence or absence of the tRNA methyltransferase DNMT2 on tRNA, rRNA and mRNA methylation levels. This identified numerous known tRNA m5C sites (Fig. 6A; Supplemental Table S5). In addition, our results also demonstrate that DNMT2 is a highly specific enzyme, as *Dnmt2* mutants specifically lost methylation at C38 in tRNA(Asp), tRNA(Gly) and tRNA(Val) (Fig. 6A). The analysis of rRNA revealed the presence of 2 novel and completely methylated cytosines in 28S rRNA, namely C3438 and C4099 (Fig. 6B). These marks were not affected in *Dnmt2* knockouts, and their sequence contexts (Supplemental Fig. S6) were identical to the m5C marks described in H. sapiens and A. thaliana rRNA (Burgess et al. 2015), consistent with evolutionary conservation of rRNA methylation patterns. In addition, we also found 2 cytosines with partial (C909, ratio equal to 0.5) and almost complete (C911, ratio equal to 0.8) methylation in mitochondrial rRNA, which is consistent with published findings (Metodiev et al. 2009). We also analyzed mRNA methylation in *Dnmt2* knockouts and could not detect any DNMT2-dependent mRNA methylation candidates (Fig. 6C). Finally, because a recent study has suggested a role of TET dioxygenases in the demethylation of *Drosophila* mRNAs (Delatte et al. 2016), we also investigated the mRNA methylation pattern of mouse ES cells that lack all three mammalian *Tet* homologues (Dawlaty et al., 2014). A comparison with mRNA methylation patterns from wild type cells again provided very little evidence for mRNA methylation (Fig. 6D). A very small number of cytosines showed a reduced non-conversion ratio in the Tet-deficient cells (Fig. 6D). However, non-conversion ratios of Tet-deficient libraries were systematically reduced by 0.004 (p-value of a two-sample t-test <2.2 x 10^-16^), which can be explained by a more efficient bisulfite deamination (Supplemental Table S2). As such, our results fail to provide any evidence for Tet-mediated demethylation of mRNAs in mouse ES cells.

**Figure 6.**
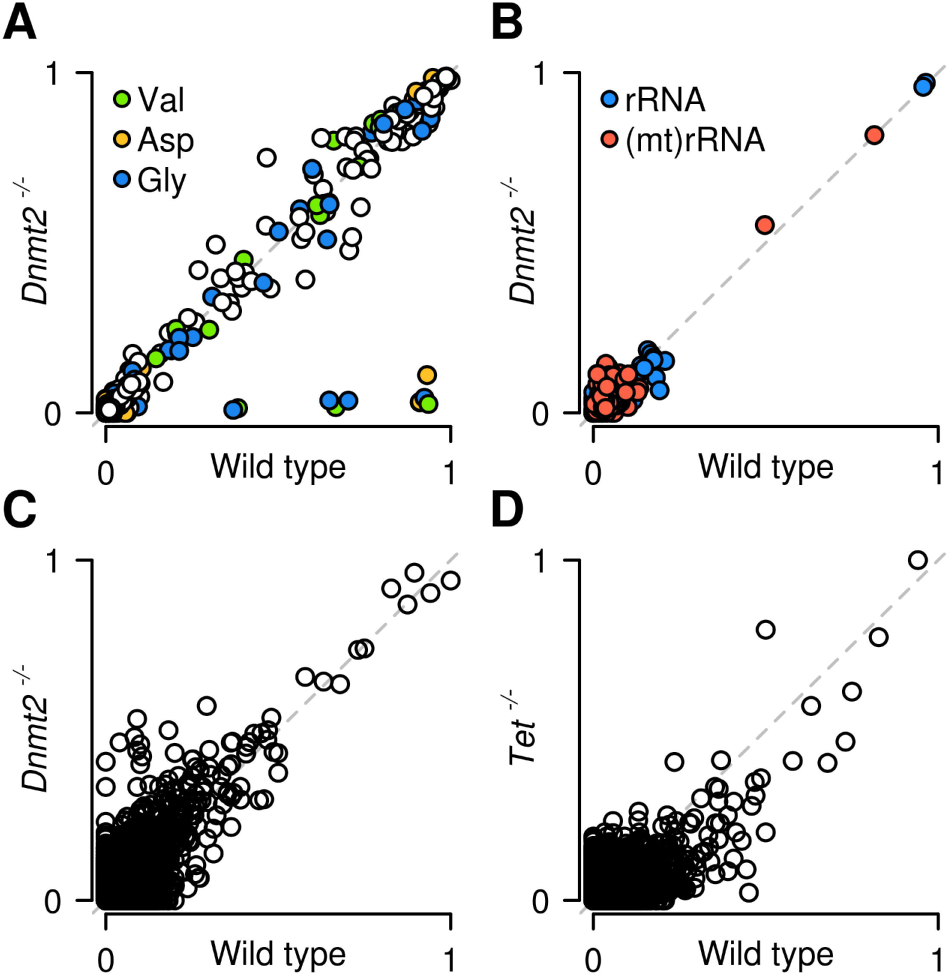
Site-specific methylation analysis by whole-transcriptome bisulfite sequencing. Scatter plots show non-converted cytosines for tRNA (*A*), rRNA (*B*), mRNA (*C*) in wildtype and *Dnmt2* knockout ES-cells. Methylation ratios are specifically reduced for C38 of tRNA(Asp), tRNA(Gly) and tRNA(Val) in *Dnmt2* knockouts. (*D*) Scatter plot for mRNA in wild type and Tet-deficient ES cells.

Together, these results comprehensively illustrate the robustness of our approach for the transcriptome-wide analysis of cytosine-5 methylation patterns at single-base resolution.

## Discussion

Previous whole-transcriptome bisulfite sequencing pipelines often relied on cutoff-based methylation calling approaches (typically coverage and non-conversion ratio >10 and >20%, respectively, Supplementary Table S6). Improved accuracy was achieved by a more stringent quality control and the removal of non-converted reads (Edelheit et al. 2013; Blanco et al. 2014), while reproducibility was ensured by the inclusion of replicates (Amort et al. 2013; Blanco et al. 2014). Most recently, it was suggested to integrate several methods derived from the DNA methylation field including generic statistical tests and a custom low-redundancy reference (Rieder et al. 2016; Amort et al. 2017). However, concerns about false positives from stochastic non-conversion events and other sources of artifacts have remained (Hussain et al. 2013a; Gilbert et al. 2016; Helm and Motorin 2017). We have now developed a pipeline for the accurate and reproducible analysis of m5C marks in whole-transcriptome bisulfite sequencing datasets. Tailored filtering addressed most sequencing and alignment artifacts, which is particularly important for the short low-complexity reads that result from bisulfite conversion. In particular, we tested for residual misalignments and discarded them if significant. Statistical modeling of bisulfite non-conversion was used to characterize and eliminate random non-conversion artifacts. Simulations confirmed a high statistical power of our pipeline, using Benjamini-Hochberg adjustment for multiple testing, or Independent Hypothesis Weighting for cytosines with a non-conversion ratio ≥20%. The inclusion of replicates allowed us to identify candidate sites that are reproducibly methylated. Also, standard errors were estimated on replicates, indicating how variable methylation is at each cytosine. The application of this pipeline on various datasets provided novel insight into the distribution of m5C in the mouse transcriptome.

A recent study detected m5C in prokaryotic mRNA, but not in yeast mRNA, suggesting that cytosine-5 mRNA methylation may be restricted to prokaryotic transcriptomes (Edelheit et al. 2013). Our results are consistent with these findings and argue against the notion that m5C is a widespread modification of coding and noncoding RNAs in mammals (Squires et al. 2012). Our results are also in agreement with earlier chromatographical studies that have failed to reveal any evidence for cytosine-5 methylation in mammalian mRNA (Desrosiers et al. 1974; Adams and Cory 1975; Salditt-Georgieff et al. 1976). Alternatively, the small amounts and high variability could indicate a rapid turnover of m5C in mRNA.

Whether the few remaining candidate sites identified in our analysis represent genuine methylation marks or reproducible deamination artifacts remains to be determined by truly orthogonal approaches, such as sequence-specific mass spectrometric analysis (Ross et al. 2016) or single-molecule real-time sequencing (Vilfan et al. 2013). It should be noted that bisulfite deamination artifacts can be caused by residual proteins that bind to nucleic acids and/or by RNA secondary structures (Supplemental Table S1), which could provide an explanation for their reproducibility.

As bisulfite sequencing cannot discriminate between m5C and hm5C (Huang et al. 2010), our results predict that hm5C also represents a very rare or absent modification in mRNA from mouse ES cells, consistent with mass spectrometry data obtained in our study and by others (Fu et al. 2014). Recent data suggesting high levels of hm5C in the *Drosophila* transcriptome (Delatte et al. 2016) may have been influenced by antibody-based detection methods. Alternatively, high levels of cytosine modification in *Drosophila* mRNA may also be a tissue-specific feature. Additional data based on direct detection methods will be required to clarify this issue.

In conclusion, our results establish whole-transcriptome bisulfite sequencing as a powerful method for a single-base resolution analysis of m5C RNA methylation patterns and suggest profound differences between the patterns of tRNA, rRNA and mRNA methylation. Our approach identified all known methylation marks in tRNA and two previously unknown but evolutionarily conserved marks in mouse 28S rRNA. Furthermore, the catalytic activity of the DNMT2 RNA methyltransferase was resolved at single-base resolution. This suggests that whole-transcriptome bisulfite sequencing can be used to profile cytosine-5 RNA methylation patterns in many experimental contexts ranging from basic biological processes to human disease.

## Methods

### Cell culture, RNA isolation and library preparation

Mouse embryonic stem cells were grown on a primary MEF feeder layer in standard medium. RNA isolation was performed with TRIzol (Ambion). 30 μg of total RNA was fractionated into a long (>200nt) and small RNA fraction (<200nt) and depleted for rRNA as indicated. Small and long fractions were DNase digested and bisulfite converted using the EZ RNA methylation kit (Zymo Research). After stepwise RNA end repair and further purification, cDNA synthesis and library preparation was carried out using the NEBNext Small RNA library Prep Set, followed by paired-end sequencing on an Illumina HiSeq 2000 platform. Further details are provided in Supplemental Methods.

### 454 (Roche) bisulfite sequencing

454 (Roche) bisulfite sequencing was performed using the EZ RNA methylation kit (Zymo Research). PCR primers are provided in Supplemental Table 7. Sequenced reads from individual amplicons were aggregated in heatmaps. Further details are provided in Supplemental Methods.

### LC-MS/MS analysis

LC-MS/MS analysis of RNA from mouse ES cells and *Drosophila* S2 cells (cultured under standard conditions) was performed as described previously (Kellner et al. 2014). Further details are provided in Supplemental Methods.

### Reference sequences

Separate references were generated for tRNA, rRNA and mRNA: (1) tRNA genomic sequences for mouse mm9 were retrieved from the genomic tRNA database. Duplicate sequences were removed. In case of bisulfite-conversion duplicates (i.e. two sequences becoming identical upon cytosine to thymine conversion), the duplicate sequence containing thymine was removed. This allows methylation detection in such sequences, even though the methylation ratio cannot be determined here. (2) Sequences for 5.8S, 18S and 28S rRNAs were retrieved from the BK000964.3 reference sequence in the NCBI nucleotide database, whereas mitochondrial rRNA sequences were obtained from Ensembl GRCm38 (release 81). We chose to keep only one variant for each main Svedberg category of rRNA (5.8S, 18S, 28S) so as to not discard reads which would otherwise align to multiple rRNA reference sequences. (3) mRNA transcript sequences were downloaded from NCBI RefSeq for mouse mm9. To ensure that reads uniquely align to mRNA, this reference was complemented with controls consisting of tRNA sequences and a comprehensive collection of non-coding RNA sequences (BK000964.3, other rRNA variants from Arb-Silva database and Ensembl, and non-coding RNA sequences from NCBI RefSeq).

### Sequence alignment

Reads were trimmed with a quality cutoff of 30 for each base (corresponding to a confidence level of 99.9%) and aligned using BSMAP (Xi and Li 2009), which allows C-T base modifications without flagging them as mismatches. We used a 3% mismatch rate, and full sequence usage in BSMAP (parameters: -s 12 -v 0.03 -g 0 -w 1000 - S 0 -p 1 -V 1 -I 1 -n 0 -r 2 -u -m 15 -x 1000). Resulting aligned reads were kept only if their length was 25 nt or more, if they aligned uniquely, and if both forward and reverse pairs were located at the same positions.

### Methylation calling

Reliable distinction between stochastic non-conversion and methylation (or other events) depends on a valid null hypothesis for the underlying distribution of stochastic non-conversion. Thus, we assessed the adequacy of the empirical null distribution formed by the non-converted cytosine counts (restricted to non-conversion lower than 0.3) to the binomial ℬ, negative binomial 𝒩ℬ or Poisson 𝒫 distributions. We estimated the parameters of each distribution and calculated the data log-likelihood using function ‘fitdistr’ in R, where the log-likelihood is denoted 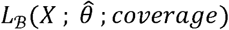, 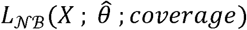 or 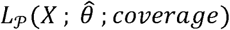 and depends on the vector containing the count of non-converted Cs at each site, *X*, on the estimated distribution parameters set, 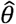, and on the coverage. Bins of 10 in coverage were used, and a minimum of 60 data points (number of cytosine positions falling within this bin) was required before applying ‘fitdistr’. Since the parameters of the binomial distribution could not be found using ‘fitdistr’, we estimated the binomial parameters 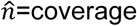 and 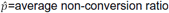 non-conversion ratio, and generated the corresponding values 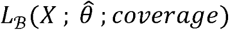. At all coverages, we obtained 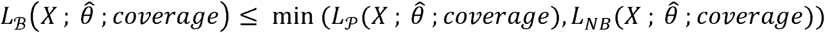, so that we considered ℬ a less useful assumption than 𝒩ℬ or 𝒫. Because 𝒫 offered a minimal and coverage-independent parameterization λ_*p*_/*coverage* (Fig. 2B; λ_*p*_; denotes the estimate of the Poisson parameter), we made the assumption that the theoretical null distribution of bisulfite non-conversion in this dataset follows 𝒫. Subsequently, we tested non-converted read counts at each specific cytosine against the null hypothesis that the counts follow 𝒫 (λ_*p*_) (Poisson exact test in R, adjustment of p-values for multiple testing with Benjamini-Hochberg method, significance level α = 0.05). A significant result was interpreted as a possibly methylated cytosine and qualified the site as a valid candidate. The adjusted p-values of cytosines which had a non-conversion ratio higher than λ_*p*_/*coverage* in each of the three replicates were combined using Fisher’s method (χ^2^ test on the statistic 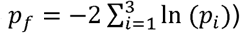. The methylation calling methods described in this paragraph are implemented in the R package BisRNA.

### Calculation of deamination rates

Deamination rates were calculated as the count of converted cytosines divided by the sum of converted and non-converted cytosines. This calculation is carried out on nuclear and mitochondrial rRNA. Known methylation sites in rRNA were removed from the calculations.

## Data Access

The sequencing data from this study have been submitted to the NCBI Gene Expression Omnibus (GEO; http://www.ncbi.nlm.nih.gov/geo/) under accession number GSE81825.

## Software availability

BisRNA is available as a source R package in the Supplemental Information and from the Comprehensive R Archive Network (https://cran.r-project.org).

## Acknowledgements

We thank Matthias Schaefer and Sebastian Bender for their contributions during early stages of this project and the DKFZ Genomics and Proteomics Core Facility for sequencing services. We also thank Günter Raddatz and Miriam Kesselmeier for critical discussion of the simulations and Michaela Frye for critical reading of the manuscript. This work was supported by grants from Landesstiftung Baden-Württemberg (Forschungsprogramm “nicht-kodierende RNAs”) to F.L. and Deutsche Forschungsgemeinschaft (Priority Programme 1784) to M.H. and F.L.

## Disclosure Declaration

The authors declare no conflicts of interest.

